# Transmembrane coupling accelerates the growth of liquid-like protein condensates

**DOI:** 10.1101/2024.11.07.622512

**Authors:** Yohan Lee, Feng Yuan, Jerry L. Cabriales, Jeanne C. Stachowiak

## Abstract

Timely and precise assembly of protein complexes on membrane surfaces is essential to the physiology of living cells. Recently, protein phase separation has been observed at cellular membranes, suggesting it may play a role in the assembly of protein complexes. Inspired by these findings, we observed that protein condensates on one side of a planar suspended membrane spontaneously colocalized with those on the opposite side. How might this phenomenon contribute to the assembly of stable transmembrane complexes? To address this question, we examined the diffusion and growth of protein condensates on both sides of membranes. Our results reveal that transmembrane coupling of protein condensates on opposite sides of the membrane slows down condensate diffusion while accelerating condensate growth. How can the rate of condensate growth increase simultaneously with a decrease in the rate of condensate diffusion? We provide insights into these seemingly contradictory observations by distinguishing between diffusion-limited and coupling-driven growth processes. While transmembrane coupling slows down diffusion, it also locally concentrates condensates within a confined area. This confinement increases the probability of condensate coalescence and thereby enhances the overall rate of growth for coupled condensates, substantially surpassing the growth rate for uncoupled condensates. These findings suggest that transmembrane coupling could play a role in the assembly of diverse membrane-bound structures by promoting the localization and growth of protein complexes on both membrane surfaces. This phenomenon could help to explain the efficient assembly of transmembrane structures in diverse cellular contexts.

**Significance:** Protein assemblies that span biological membranes are critical to cellular physiology. In the past decade, liquid-like protein condensates, which are flexible, multivalent protein assemblies, have been discovered on diverse membrane surfaces. Recently, we observed that protein condensates on opposite sides of a membrane spontaneously colocalize to form coupled, transmembrane complexes. Interestingly, while transmembrane coupling slows down the diffusion of membrane-bound condensates, it substantially accelerates their growth by strongly localizing interactions between them. These findings suggest that transmembrane coupling of protein condensates may play a role in promoting the robust assembly of membrane-bound protein complexes in crowded, complex cellular environments.

## Introduction

Dynamic protein complexes on membrane surfaces, including focal adhesions,^1,2^ immunological synapses,^3–5^ cell-cell junctions,^6,7^ and endocytic vesicles,^8,9^ rely on the precise assembly of proteins at specific membrane sites. These structures arise from the sequential recruitment of their constituent proteins to both surfaces of the plasma membrane. The spatial and temporal regulation of protein localization is key to forming these assemblies, yet the mechanism underlying their dynamic assembly remains incompletely understood. A critical question is how functional transmembrane complexes assemble efficiently in the crowded, complex environment of cellular membranes.^10^ Recently, liquid-like protein condensates, flexible, multivalent protein assemblies, have been discovered on membrane surfaces.^7,8,11,12^

Can the formation of liquid-like condensates on the membrane drive the assembly of transmembrane complexes? Previous work has investigated the dynamics of condensates composed of proteins, nucleic acids, or polymers on the surface of lipid vesicles^13–15^ and supported lipid bilayers.^11,16–19^ However, these studies were limited to condensates on only one side of the membrane. In recent work, we addressed this gap by introducing suspended planar lipid membranes as a unique platform to study protein phase separation on both membrane surfaces.^20^ Using this system, we observed a transmembrane coupling phenomenon in which liquid-like protein condensates on opposite sides of a membrane spontaneously colocalized with one another. We found that protein condensates locally reorganized lipid membranes, reducing lipid mobility and resulting in reduced lipid entropy. As a result, transmembrane coupling of protein condensates on opposite sides of the membrane became thermodynamically favorable, as it minimized the loss in lipid entropy, thereby maximizing the overall entropy of the coupled system.^20^

Here, we investigate the impact of transmembrane coupling on the dynamics of liquid-like protein condensates. Surprisingly, our results reveal that, while transmembrane coupling slows the diffusion of condensates, it simultaneously accelerates their coarsening. We resolve this paradox by distinguishing between diffusion-limited and coupling-driven coarsening processes. Specifically, our findings suggest that transmembrane coupling facilitates the concentration of condensates within a confined protein region, increasing the likelihood of condensate contact and thereby accelerating overall coarsening. These findings provide new insights into how transmembrane coupling of protein condensates might contribute to the assembly of transmembrane protein complexes.

## Results

### Membrane-bound liquid-like protein condensates exhibit Brownian diffusion

We created suspended planar lipid membranes within the hexagonal holes of transmission electron microscopy (TEM) grids, as described previously.^20^ Each grid contained 150 holes, each with a diameter of approximately 100 μm, enabling visualization of multiple independent membrane surfaces per field of view using fluorescence confocal microscopy (Figure 1a). To investigate the dynamics of protein condensates on planar lipid membranes, the RGG domain of the LAF-1 protein (RGG) was used, as its condensation by protein phase separation is well-established.^21^ We added N-terminal histidine-tagged RGG (his-RGG), labeled with Atto 488, at a concentration of 1 μM to suspended planar membranes containing DGS-Ni-NTA lipids (15-25 mol%). Here, binding of his-RGG to the membrane was achieved through interactions between histidine and Ni-NTA. In our initial experiments, we sought to examine condensate dynamics on only one side of the membrane. Therefore, we placed the TEM grid directly against a glass coverslip so that proteins only bound significantly to the top surface (Figure 1a). Importantly, RGG is highly soluble in aqueous buffers, such that phase separation in solution only occurs at concentrations greater than 10 μM, substantially above the concentration used in our experiments with membranes.^21,22^ In this way, we could differentiate protein phase separation on membranes from phase separation in the surrounding solution, which was not observed under our experimental conditions. Initially, the fluorescence intensity in the protein channel was homogeneous over the surface of the membrane, indicating uniform protein binding. However, brighter and dimmer regions began to coexist after a few minutes, indicating phase separation of the membrane-bound protein layer into protein-rich (brighter) and protein-depleted (dimmer) phases (Figure 1b). The membrane-bound protein-rich condensates had round shapes, merging and re-rounding upon contact, suggesting that they had liquid-like properties (Figure 1c).

**Figure 1.**
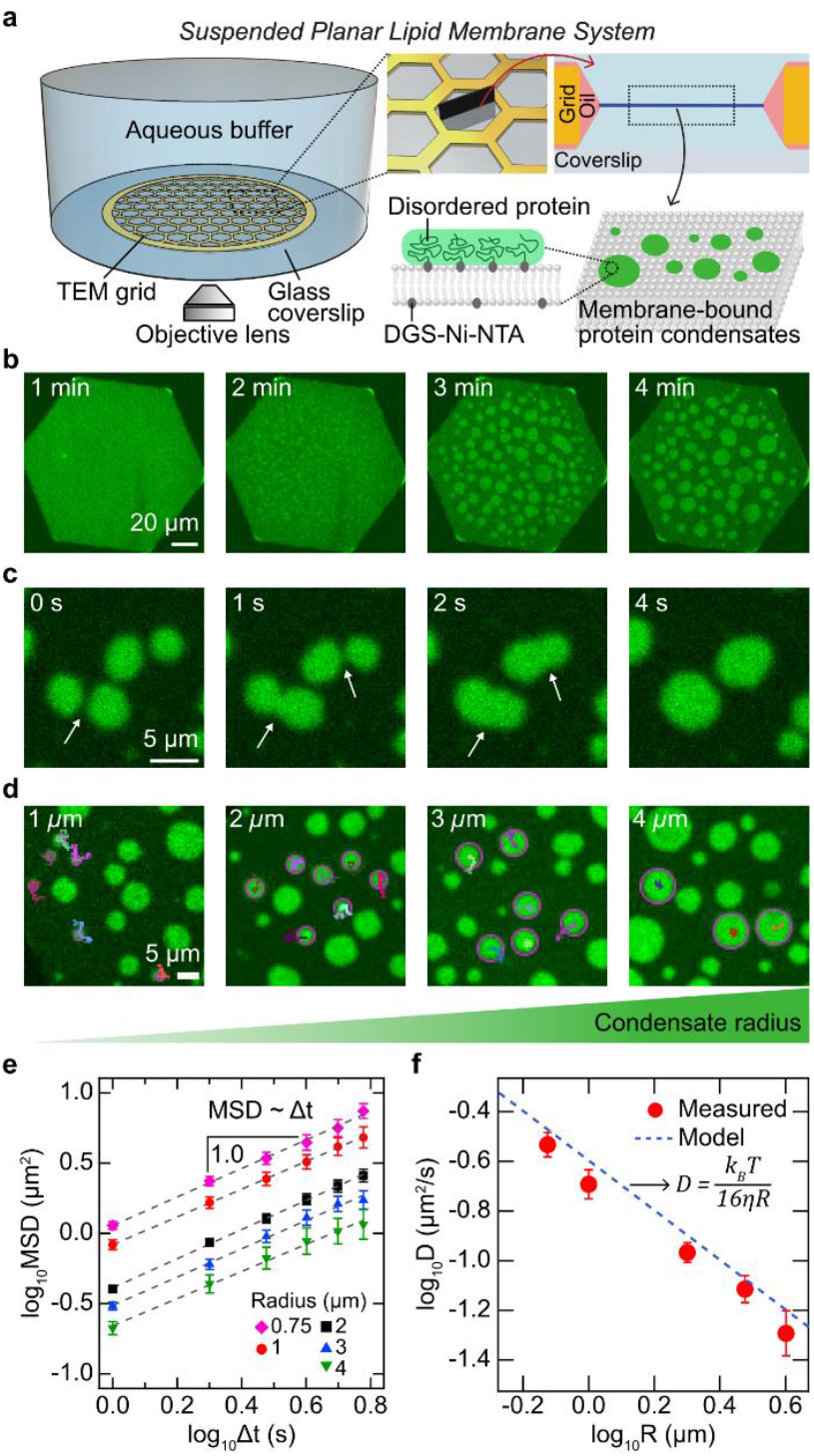
Diffusion of RGG protein condensates bound on the top surface of the membrane exhibits Brownian motion. (a) Schematic of the suspended planar membrane system where suspended membranes span each hexagonal hole of a transmission electron microscopy (TEM) grid. Histidine-tagged proteins with disordered regions, in this case, RGG, bind to the membrane through histidine-nickel interactions and form protein condensates via spontaneous protein phase separation. (b) Representative images showing protein phase separation on the membrane at specified elapsed time. Scale bar, 20 μm. (c) Merging and re-rounding of membrane-bound protein condensates. White arrows indicate merging spots. Scale bar, 5 μm. (d) Protein condensates were tracked over time according to their radii. Scale bar, 5 μm. Tracked condensates (magenta circles) with trajectories are shown. (e) MSDs over time lags (Δt) for RGG condensates with a radius of 0.75 μm (magenta diamonds, n = 38), 1 μm (red circles, n = 24), 2 μm (black squares, n = 97), 3 μm (blue triangles, n = 45), and 4 μm (green inverted triangles, n = 21). The dashed lines represent a linear fit: log_10_MSD = b + α·log_10_Δt. For all MSDs, the fitted slopes were 1.0 (α = 1), confirming MSD ∼ Δt, a characteristic of Brownian motion. (f) Diffusivity (D) for each condensate radius (R) was obtained as 4·D = 10^b^ using the fitted results in (e): MSD = 10^b^ Δt^α^. The dashed blue line represents a model relating D and R for large domain size, that is, D = k_B_T / (16ηR), where k_B_ is the Boltzmann constant, T is the temperature (293 K), and η is the water viscosity (1.0 cP). Error bars represent standard errors. Membrane composition: 85 mol% DOPC, 15 mol% DGS-Ni-NTA, and 0.5 mol% Texas Red-DHPE. Buffer: 25 mM HEPES, 100 mM NaCl, pH 7.4. 1 μM of his-RGG labeled with Atto 488 was used.

First, we sought to examine the diffusion of protein condensates on membranes composed primarily of 1,2-dioleoyl-sn-glycero-3-phosphocholine (DOPC) lipids (full composition in Figure 1 caption) by tracking their position over time and calculating the mean squared displacement (MSD) over time lags (Δt) (Figure 1d, Movie S1). Generally, MSD has a power-law relationship with time, expressed as MSD ∼ Δt^α^, where α represents the diffusion exponent. In the case of normal diffusion, which is governed by Brownian motion, the diffusion exponent is 1 (MSD ∼ Δt). Conversely, anomalous diffusion is characterized by a diffusion exponent that deviates from 1, greater than 1 for superdiffusion, and less than 1 for subdiffusion. We obtained MSDs as a function of Δt for condensates with radii ranging from 0.75 μm to 4 μm, in which all MSDs had a diffusion exponent of 1, indicating normal diffusion by Brownian motion (Figure 1e).

Over the past several decades, several groups have studied Brownian diffusion of domains on membrane surfaces. The Saffman-Delbrück model proposes that the diffusivity (D) of a domain is proportional to the logarithm of its radius (R), such that D ∼ ln(1/R).^23–27^ A key parameter in this model is the Saffman-Delbrück length (L_SD_), defined as L_SD_ = η_2D_/2η_3D_, where η_2D_ represents the membrane viscosity, and η_3D_ is the bulk viscosity of the surrounding medium, typically water. The Saffman-Delbrück model holds for small domains, where R < 0.1×L_SD_,^24^ which corresponds to R < 200 nm in our system, using η_2D_ ∼ 4 × 10^−9^ Pa·s·m (for DOPC membrane) and η_3D_ ∼ 1 × 10^−3^ Pa·s.^28^ For larger domains, the diffusivity can be approximated as D = k_B_T / 16η_3D_R, where k_B_ is the Boltzmann constant, and T is the temperature, indicating that diffusion is dominated by the bulk fluid surrounding the membrane.^24,29–31^ Since the radii of membrane-bound protein condensates in our system are several micrometers, we expect their diffusion to follow the latter model.

We extracted the diffusivity from the MSDs and plotted it against the condensate radius, and determined that diffusivity was inversely proportional to radius, as expected (Figure 1f). We obtained similar results by examining the diffusion of membrane-bound condensates composed of another highly studied condensate-forming protein, the low complexity domain of the fused in sarcoma protein (FUS LC) (Figure S1).^32,33^ Phase separation of FUS LC is driven by pi-pi stacking among tyrosine residues, while phase separation of RGG primarily relies on electrostatic interactions.^34^ Despite the different protein sequences and condensation mechanisms of RGG and FUS LC, the similar diffusion behavior of the resulting condensates suggests that this behavior is independent of the protein identity and specific phase separation mechanism. Collectively, we analyzed the diffusion of protein condensates bound to one side of a suspended lipid bilayer and found that they undergo random, normal diffusion.

### Uncoupled condensates experience diffusion-limited coarsening

As condensates diffused over the membrane surface, they frequently collided, resulting in merging and re-rounding over a timescale of seconds, leading to an increase in condensate size over time (Figure 1c and Figure 2a). We next examined the dynamics of this coarsening behavior. By tracking pairs of colliding condensates, we confirmed that when two condensates merged, their surface area was conserved such that the area of the final condensate was equal to the sum of the areas of the original condensates (Figure 2b,c).

**Figure 2.**
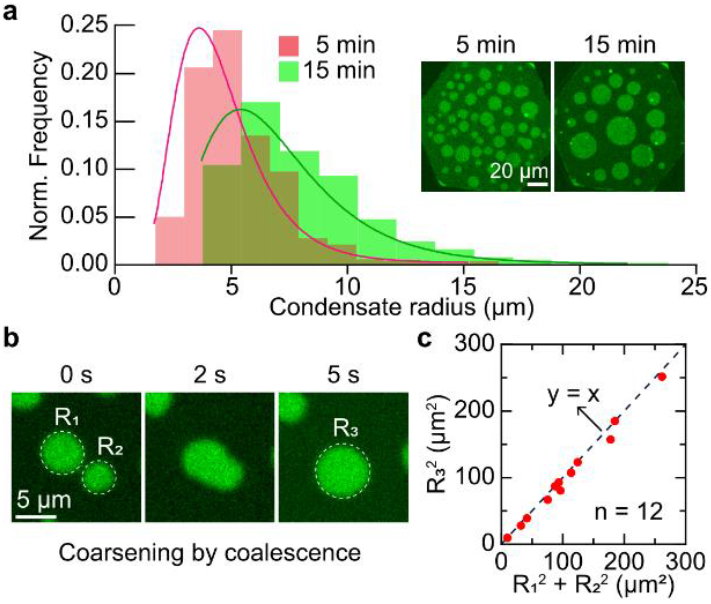
The coarsening of protein condensates on a one-sided membrane system. (a) Histogram showing the fraction of condensates depending on their radii after 5 minutes (n = 478) and 15 minutes (n = 210). Solid curves are Log-normal fitting results. (Right) Representative images of membrane-bound protein condensates at specified elapsed time. Scale bar, 20 μm (b) Membrane-bound protein condensates merging and re-rounding upon contact. R_1_ and R_2_ represent the radius of each condensate, marked by dashed circles, before coalescence and R_3_ after coalescence. Scale bar, 5 μm (c) Area conservation holds over multiple coalescence events (n = 12). The dashed line represents the y = x line.

To further probe the coarsening dynamics of membrane-bound RGG condensates, we monitored their area over time. The increase in area over time followed the power law A ∼ t^β^, where A, t, and β represent area, time, and the growth exponent, respectively. It is well established that a growth exponent of 2/3 holds for the particles undergoing a normal diffusion following D ∼ 1/R. This is because the spacing (l) between domains is proportional to their radius, l ∼ R. For these domains to coarsen by coalescence, they must travel an average distance l, which requires a time, t ∼ l^2^/D ∼ R^2^/D, resulting in A ∼ R^2^ ∼ t^2/3^. This growth exponent has been observed in diverse biological systems, including lipid membranes with phase-separated lipid domains,^31^ and protein droplets, both in vitro and in living cells.^35,36^

We first examined the coarsening dynamics of RGG condensates on membranes composed primarily of DOPC lipids, the same as above (full composition in Figure 3 caption). Our analysis revealed a growth exponent of 2/3, consistent with normal diffusion of the condensates (Figure 3a,d). To determine whether membrane composition impacted this result, we conducted the same analysis on membranes mainly composed of 1-stearoyl-2-oleoyl-sn-glycero-3-phosphocholine (SOPC) lipids (full composition in Figure 3 caption). While DOPC has two unsaturated oleic acid groups attached to a phosphocholine head group, SOPC has an unsaturated oleic acid group and a saturated stearic acid group as hydrophobic tails. Due to the different tail group chemistries of the two lipids, the melting temperature (T_m_) of SOPC (6 °C) is higher than that of DOPC (−17 °C). Further, SOPC-enriched membranes have about twice the membrane viscosity than DOPC-enriched membranes.^28^ We observed the same growth exponent of 2/3 for RGG condensates on the SOPC-enriched membranes (Figure 3b,e). These results suggest that the coarsening of protein-rich domains is not a strong function of membrane viscosity, as expected for large domains whose diffusion is limited by the viscosity of bulk solution rather than membrane viscosity.

**Figure 3.**
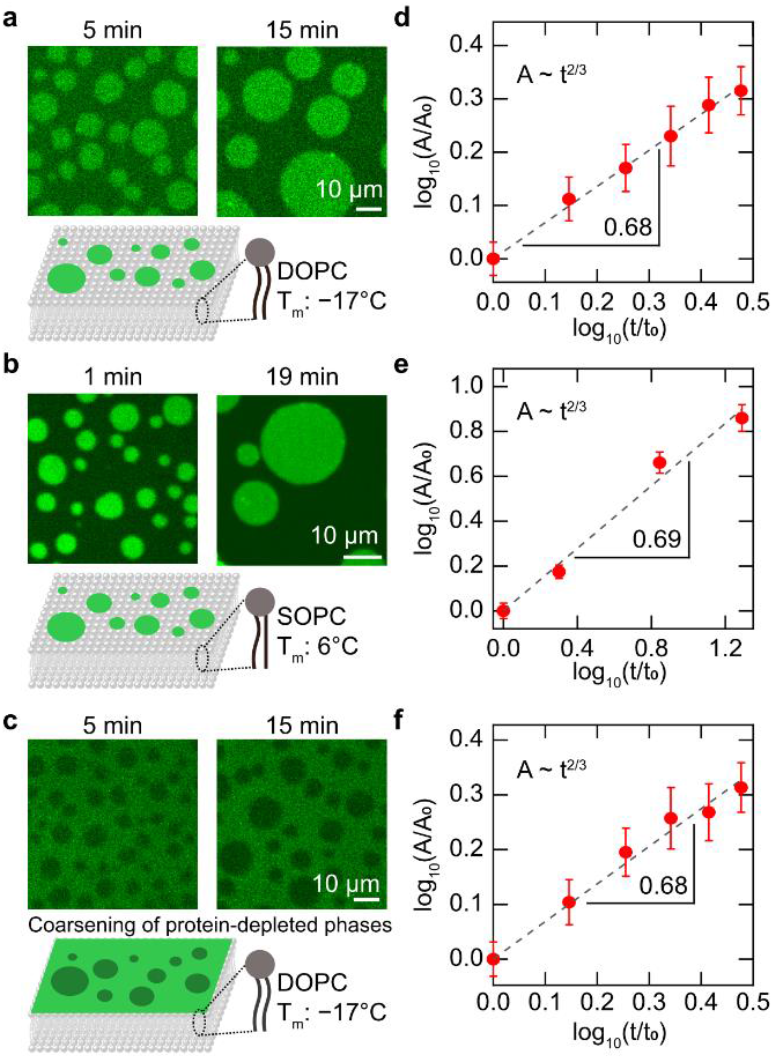
Protein condensates on a one-sided membrane system coarsen with the growth exponent of 2/3. (a-c) Snapshots showing membrane-bound RGG condensates (a, b) and RGG-depleted regions dispersed within a continuous RGG-enriched region (c) at the indicated elapsed times after protein addition. The major component of the lipid membrane was DOPC with a melting temperature (T_m_) of −17°C (a, c) and SOPC with T_m_ of 6°C (b). (d-f) Average area of RGG condensates (d, e) and RGG-depleted regions (f) were plotted over time, after normalizing by the area A_0_ of each condensate/void at the specified t_0_. The dashed line represents a linear fit: log_10_(A/A_0_) = β·log_10_(t/t_0_), with β being the growth exponent. The obtained β was 0.68 (d), 0.69 (e), and 0.68 (f), all confirming A ∼ t^2/3^. Error bars represent standard errors from analyzing 20-113 protein condensates or protein-depleted regions per time from at least three independent membranes. Scale bars, 10 μm. Membrane composition: 75-85 mol% DOPC (or SOPC), 15-25 mol% DGS-Ni-NTA, and 0.5 mol% Texas Red-DHPE. Buffer: 25 mM HEPES, 100 mM NaCl, pH 7.4. 1 μM of his-RGG labeled with Atto 488 was used.

Owing to multivalent interactions between the proteins within condensates, the protein-enriched phase is likely to be more viscous than the protein-depleted phase. In the above coarsening events, the more viscous, protein-enriched phases diffused and coarsened over time, surrounded by a continuous protein-depleted phase (Figure 3a,b). In contrast, when the concentration of DGS-Ni-NTA lipids was high (25 mol%) with 100 mM of NaCl concentration—factors that facilitate RGG phase separation—we could observe a continuous protein-enriched phase with dispersed protein-depleted phase regions (Figure 3c). Here, we might expect the protein-depleted regions to coarsen more slowly than the protein-enriched regions, owing to the likely higher viscosity of the protein-enriched continuous phase. Nevertheless, the same growth exponent of 2/3 was observed during the coarsening of the protein-depleted regions on the DOPC-enriched membrane (Figure 3f). These findings further suggest that the coarsening of membrane-bound protein condensates is governed by normal diffusion, such that it is largely insensitive to membrane viscosity.

### Transmembrane coupling slows diffusion of membrane-bound condensates

We next asked how the diffusion of condensates changes when they form on both surfaces of the membrane. By introducing thin spacers between the TEM grid and the coverslip, we enabled proteins to access both sides of the membrane (Figure 4a). Under this condition, we observed transmembrane condensate coupling, where protein condensates on opposite sides of the membrane spontaneously colocalized.^20^ These coupling events produced three distinct levels of fluorescence intensity: brightest, medium-bright, and dimmest (Figure 4b), compared to only two levels of fluorescence intensity when condensates form on a single side of the membrane (Figure 1b-d, Figure 2b, and Figure 3). Notably, the intensity difference between the dimmest and medium-bright regions was comparable to that between the medium-bright and brightest regions. This observation suggests that the brightest regions indicate protein condensates on both membrane surfaces, medium-bright regions indicate condensates on only one surface of the membrane, and dimmest regions indicate the absence of condensates on both membrane surfaces (Figure 4c). The space between the bottom membrane surface and the glass coverslip was limited to about 50 μm to meet the working distance requirements of the high magnification microscope objective, somewhat slowing diffusion of proteins to the bottom side of the membrane. As a result, protein condensates formed first on the top surface of the membrane, which was more accessible to protein in solution, followed by the gradual appearance of protein condensates on the bottom surface of the membrane. For this reason, the condensates on the bottom surface of the membrane were generally smaller than those on the top surface, especially in the early moments of our experiments. Also, regions of the brightest intensity were often surrounded by regions of medium intensity (Figure 4b). This observation suggests that smaller protein condensates on the bottom side of the membrane were coupled to a larger condensate on the top membrane surface (Figure 4c cartoon).

**Figure 4.**
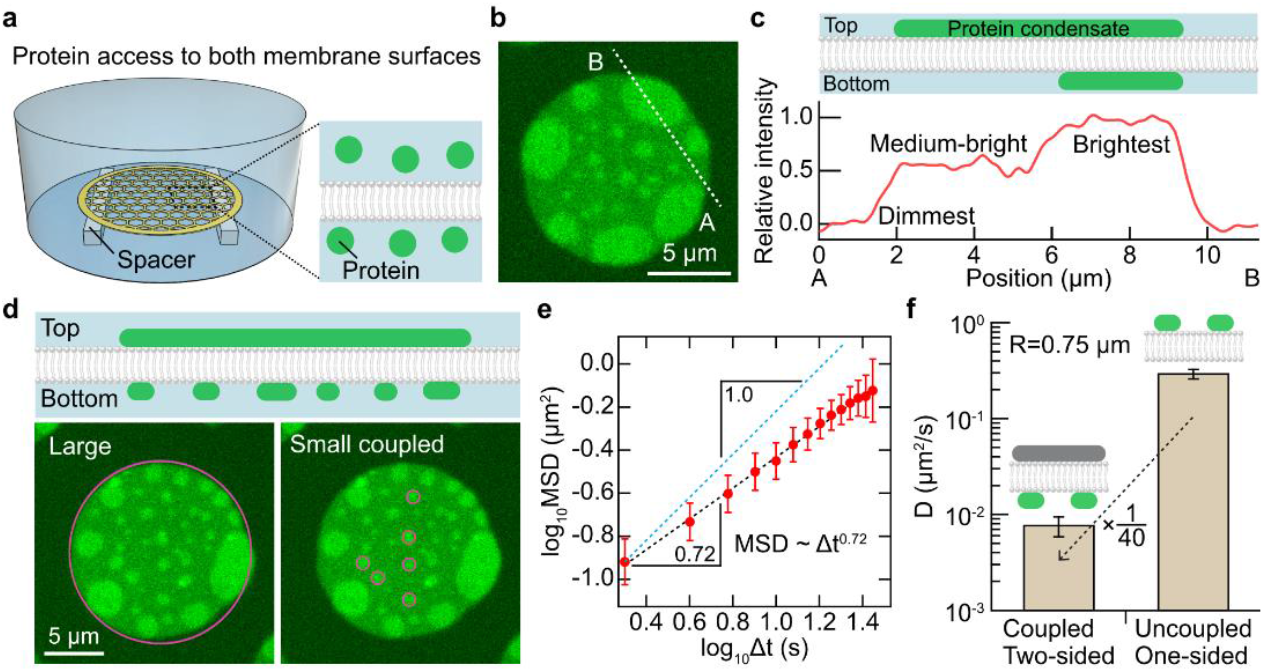
Coupled condensates exhibit subdiffusion. (a) Schematic of the suspended planar membrane system with spacers between the TEM grid and the glass coverslip where proteins are allowed to access both sides of the membrane. (b) Representative microscopic image of protein regions with three different brightnesses: dimmest, medium-bright, and brightest regions. (c) Cartoon of cross-section of the membrane (Top) and relative intensity profile (Bottom) along the dotted line (from A to B) in (b), where regions with relative intensity of 0, 0.5, and 1.0 correspond to the dimmest, medium-bright, and brightest regions, respectively. Relative intensity (I_R_) was defined as *I*_*R*_ *= (I – I*_*D*_*)/(I*_*B*_ *– I*_*D*_), where I, I_D_, and I_B_ indicate the fluorescence intensity of the region of interest, the intensity of the dimmest region, and the intensity of the brightest region, respectively. Due to limited space between the bottom membrane surface and the glass coverslip, protein binding to the bottom surface is slower than that to the top surface. Therefore, condensates on the top membrane surface are likely larger than condensates on the bottom. (d) (Top) Cartoon of a cross-section of the protein-bound membrane. (Bottom) Images highlighting a large condensate on the top membrane surface (left) and the small brightest condensates (R = 0.75 μm) on the bottom membrane surface coupled to the large condensate (right). (e) MSD over time lags (Δt) for small coupled condensates (n = 20). The dashed black line represents a linear fit: log_10_MSD = b + α·log_10_Δt. The fitted slope (α) was 0.72, indicating subdiffusion with the MSD ∼ Δt^0.72^. The dashed blue line is a reference with the slope of 1. (f) Diffusivity of condensates with radii of 0.75 μm that are coupled and uncoupled, which was obtained by the relationship MSD = 4DΔt, showing 40 times difference. The cartoon above each bar graph indicates condensates of interest colored green. Error bars represent standard errors. Membrane composition: 85 mol% DOPC, 15 mol% DGS-Ni-NTA, and 0.5 mol% Texas Red-DHPE. Buffer: 25 mM HEPES, 100 mM NaCl, pH 7.4. 1 μM of his-RGG labeled with Atto 488 was used.

We next asked how transmembrane coupling impacted the diffusion of protein condensates. For this purpose, we analyzed the diffusion of small protein condensates on the bottom side of the membrane coupled to a large protein condensate on the top side of the membrane (Figure 4d cartoon). Notably, diffusion of the large condensate on the top membrane surface influenced the motion of small, coupled condensates on the bottom membrane surface (Movie S2). To isolate the diffusion of the small condensates relative to the large condensate, we first tracked the positions of both large and small condensates. Then, we calculated the relative position of the small condensates by subtracting the position of the large condensate from that of the small ones. Using these relative positions, we computed the MSD of the small, coupled condensates over different time lags. This analysis revealed a diffusion exponent of 0.72, indicating subdiffusion (Figure 4e). We further compared the diffusivity of protein condensates with the same radius under uncoupled (one-sided, described in Figure 1) and coupled (two-sided) conditions. The diffusivity of the coupled condensates was found to be nearly 40 times lower than that of the uncoupled condensates (Figure 4f).

Why do smaller, coupled condensates exhibit reduced diffusivity? Previous work has suggested that protein condensates increase the packing of membrane lipids.^37^ Additionally, our earlier work demonstrated that protein condensates decreased the mobility of lipids beneath them.^20^ Taken together, these findings suggest that protein condensates on the top membrane surface induce reorganization of the underlying lipids, leading to a local decrease in membrane mobility. As a result, smaller condensates on the bottom surface, coupled to a large condensate on the top, may undergo subdiffusion with reduced diffusivity, which likely impacts their coarsening dynamics. Notably, these results suggest that once condensates become coupled, the viscosity of the resulting protein-lipid composite region is large enough to impact condensate diffusion. These results are in contrast to uncoupled condensates, for which diffusion and coarsening were largely independent of membrane viscosity (Figure 1f and Figure 3). We next examined how the coarsening of coupled condensates is impacted by their slow diffusion in two-sided membrane systems.

### Transmembrane coupling accelerates coarsening of membrane-bound condensates

We observed that the extent of coupling between condensates on opposite sides of the membrane increased over time, leading to an increase in the area of the coupled regions, (Figure 5a, Movie S3, brightest regions). In movies of this coupling process, it appears that coupling increases owing to small, uncoupled condensates, presumably on the bottom surface of the membrane, crossing paths with large condensates, presumably on the top side of the membrane. During these events, the brightness of the overlapped region increased while its area remained the same (Figure 5b).

**Figure 5.**
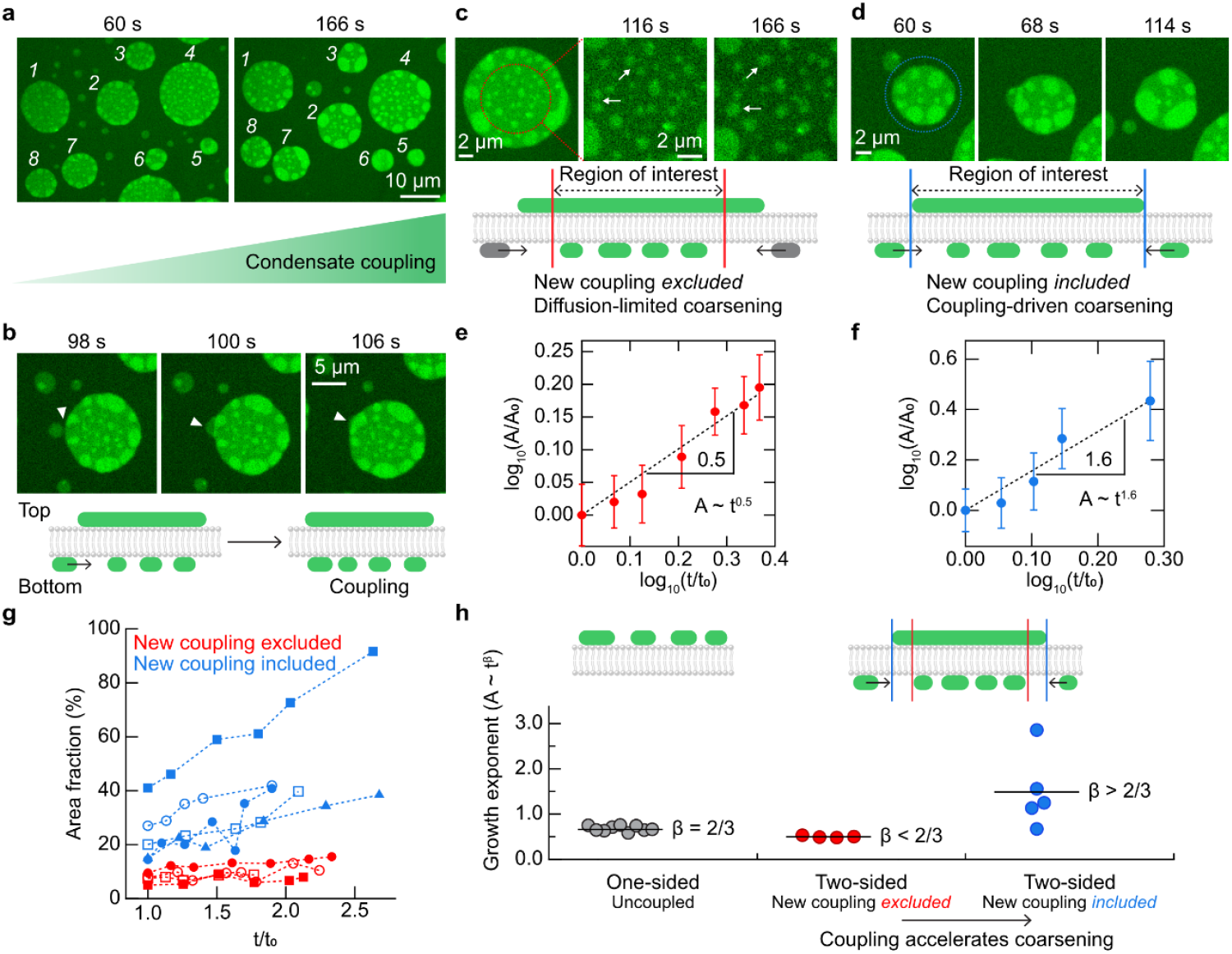
Coarsening dynamics of RGG condensates in two-sided membrane systems. (a) Images showing an increase in the extent of transmembrane condensate coupling over time. Each number in white indicates an identical protein composite region in both images. (b) Images showing a coupling process where a small condensate on the bottom side of the membrane and a large condensate on the top cross paths. Brightness increases in the overlapped region without an increase in condensate size, confirming a coupling process described in the cartoon below. (c-f) Coarsening of coupled condensates (brightest) within an inner circle of a protein composite region that excludes coupled condensate at the periphery (c, e) or within an entire protein composite region (d, f). The area of inner circles accounted for 30-40% of the entire protein region area. The average area of coupled condensates over time within the region of interest in (c) is shown in (e), and in (d) shown in (f), after normalizing by the area A_0_ of each condensate at the specified t_0_ of 60 s. The dashed line represents a linear fit: log_10_(A/A_0_) = β·log_10_(t/t_0_), with β being the growth exponent. The obtained β was 0.5 (d) and 1.6 (f). Error bars represent standard errors. (g) Area fraction of coupled condensates over time within the region of interest excluding new coupling (red) or including new coupling (blue). (h) Growth exponent from coarsening of protein condensates. For uncoupled condensates in a one-sided system, β of 2/3 was observed (gray). For coupled condensates in a two-sided system, β smaller than 2/3 was observed when new coupling was excluded (red), and β greater than 2/3 was obtained when new coupling was included (blue).

To investigate coarsening dynamics in two-sided systems, we selected several membrane-bound protein composite regions, each composed of a large condensate on one side of the membrane and multiple smaller coupled condensates on the other side of the membrane (Figure 5a). We then measured the average area of the coupled regions over time. We observed two distinct modes of coarsening in these protein composite regions: diffusion-limited and coupling-driven coarsening. The diffusion-limited coarsening occurs within the center of a protein composite region, where small condensates cross paths with one another and fuse together. This type of coarsening primarily depends on the diffusion and coalescence of small, coupled condensates, a process which is likely to be slowed down by low membrane mobility in the composite regions. Meanwhile, the coupling-driven coarsening is driven by interactions between newly coupled condensates near the periphery of the protein composite regions.

To examine these two modes of coarsening, we conducted separate analyses within two different regions of interest: (i) the center of the selected protein composite region, excluding the coupled condensates at the periphery (Figure 5c); and (ii) the entire region of the selected protein composite region, where the newly coupled regions are included (Figure 5d). For the diffusion-limited coarsening, if the subdiffusion of coupled condensates on the bottom surface slows down their coarsening, the growth exponent should be smaller than 2/3. As expected, we obtained the growth exponent of ∼ 0.5 for these condensates (Figure 5e). How will the coarsening be affected if the newly coupled regions at the edge of the larger condensate are considered? Surprisingly, for the coupling-driven coarsening, the average growth exponent was 1.6, more than twice the growth exponent of 2/3 for coarsening of uncoupled condensates (Figure 5f).

How does inclusion of the newly coupled regions impact the coarsening process? In the case of diffusion-limited coarsening with the newly coupled regions excluded, the area fraction of coupled regions remains low (10% on average) and constant (red markers in Figure 5g). In contrast, for the coupling-driven coarsening where newly coupled regions are included, the area fraction increases over time, approaching 100% (blue markers in Figure 5g). From these observations it appears that the continuous influx of new condensates via coupling into a confined protein composite region increases the area fraction of coupled regions, which, in turn, raises the probability of condensate contact, resulting in more rapid coarsening.

## Discussion

Collectively, our results demonstrate that transmembrane coupling of protein condensates accelerates condensate growth. Specifically, we show that the presence of condensates on both sides of the membrane facilitates localization and coarsening of condensates within confined membrane-bound regions. On the one hand, transmembrane condensate coupling slows the diffusion of coupled condensates, likely due to cross-linking of lipids by the proteins within the condensate. On the other hand, coupling continuously concentrates protein condensates in a confined region, which increases contact among condensates, accelerating coarsening. In cells, spatiotemporal control of protein clustering on both membrane surfaces is critical for diverse membrane-bound structures. In this context, transmembrane coupling of protein condensates may help to accelerate dynamic assembly of membrane complexes.

We have investigated the diffusion and coarsening of liquid-like protein condensates on reconstituted suspended planar membranes. We demonstrate that membrane-bound condensates undergo Brownian diffusion, following the relationship D ∼ 1/R. As a consequence of this normal diffusion, the coarsening of membrane-bound condensates follows the classic relationship A ∼ t^2/3^. Notably, this growth exponent might be different for small, diffraction-limited condensates, where the influence of membrane viscosity could dominate that of bulk fluid viscosity. In contrast, when condensates couple across the membrane bilayer, their diffusivity is reduced, likely owing to cross-linking of lipids by condensates. This reduction in diffusivity results in fewer collisions among coupled condensates, slowing coarsening within coupled regions. However, when we also considered the contribution of new condensates colliding with coupled regions, the overall rate of coarsening increased, even exceeding the growth rate of uncoupled condensates only on one side of the membrane (Figure 5h).

How can these phenomena contribute to the spatiotemporal regulation of transmembrane protein complexes in crowded cellular environments, where protein diffusion is restricted and slow?^10^ Our results suggest that once a protein condensate forms on one side of the membrane, it serves as a platform for accelerating the growth of coupled condensates on the opposite side of the membrane. Transmembrane condensate coupling is thermodynamically favorable because it minimizes the loss in lipid entropy resulting from protein condensation on membrane surfaces, thereby maximizing the overall entropy of the coupled system.^20^ By harnessing this thermodynamic driving force, transmembrane condensate coupling could work together with lipid phase separation^38^ and depletion-attraction effects^39^ to promote the assembly of functional transmembrane structures in complex and crowded cellular environments.

## Materials and Methods

### Materials

1,2-dioleoyl-sn-glycero-3-phosphocholine (DOPC), 1-stearoyl-2-oleoyl-sn-glycero-3-phosphocholine (SOPC), and 1,2-dioleoyl-sn-glycero-3-[(N-(5-amino-1-carboxypentyl)iminodiacetic acid)succinyl] nickel salt (DGS-Ni-NTA) were purchased from Avanti Polar Lipids. Texas Red 1,2-dihexadecanoyl-sn-glycero-3-phosphoethanolamine triethylammonium salt (Texas Red-DHPE), and 4-(2-hydroxyethyl)-1-piperazineethanesulfonic acid (HEPES) were purchased from Thermo Fisher Scientific. Sodium chloride, sodium tetraborate, hexadecane, silicone oil AR 20, poly-L-lysine MW 15,000-30,000 (PLL), Atto 488 NHS ester was purchased from Sigma-Aldrich. Amine-reactive PEG (mPEG–succinimidyl valerate, MW 5000) was purchased from Laysan Bio.

### Plasmids

The plasmid for RGG (pET-RGG) was a gift from Matthew Good, Daniel Hammer, and Benjamin Schuster (Addgene plasmid #124929; https://www.addgene.org/124929).^22^ The plasmid for FUS LC (RP1B FUS 1-163) was a gift from Nicolas Fawzi (Addgene plasmid #127192; https://www.addgene.org/127192).^33^

### Protein expression and purification

Expression and purification of the RGG domain of LAF-1 protein (RGG) was performed as described previously.^22^ Briefly, E. Coli BL21(DE3) competent cells were transformed with a plasmid encoding the RGG domain of LAF-1. After transformation, cells were grown in 1 L of 2xYT media for 3-4 hours at 37 °C while shaking at 220 rpm until the optical density at 600 nm (OD 600) of the media became reached 0.8, followed by overnight expression induced with 0.5 mM of isopropyl β-D-1-thiogalactopyranoside (IPTG) at 18 °C while shaking at 220 rpm. Pellets of cells expressing RGG were harvested through centrifugation at 4 °C. Pellets were resuspended in 40 mL buffer containing 20 mM Tris, 500 mM NaCl, 20 mM imidazole, 1% Triton X-100, and one EDTA-free protease inhibitor tablet (Sigma Aldrich) at pH 7.5, and lysed by sonication on ice. To prevent the formation of RGG condensates, all of the following steps were done at room temperature. The cell lysate was clarified by centrifugation at 15,000 x g for 30 min and then incubated with Ni-NTA resin (G Biosciences, USA) for 1 hour to achieve binding. Protein-bound Ni-NTA resin was settled in a glass column and washed with a buffer containing 20 mM Tris, 500 mM NaCl, and 20 mM imidazole at pH 7.5. The bound proteins were eluted from the Ni-NTA resin with a buffer containing 20 mM Tris, 500 mM NaCl, and 500 mM imidazole at pH 7.5. Purified proteins were then exchanged into the storage buffer (20 mM Tris, 500 mM NaCl, pH 7.5). Small aliquots of the protein were flash-frozen using liquid nitrogen and stored at −80 °C.

Expression and purification of the low-complexity domain of fused in sarcoma (FUS LC) was carried out according to previous reports.^32,40^ In brief, FUS LC was overexpressed in E. Coli BL21(DE3) cells. Cells were grown for 4 hours at 37 °C while shaking at 220 rpm until OD 600 reached 0.8, followed by expression induced with 1 mM of IPTG at 37 °C for 3 hours. Pellets of cells expressing FUS LC were collected and lysed by sonication on ice in 40 mL lysis buffer containing 500 mM Tris pH 8.0, 5 mM EDTA, 5% glycerol, 10 mM β-mercaptoethanol, 1 mM phenylmethylsulfonyl fluoride (PMSF), 1% Triton X-100 and one EDTA-free protease inhibitor tablet (Sigma Aldrich). The cell lysates were centrifuged at 125,000 x g for 40 min at 4°C. Unlike RGG, FUS LC resided in the insoluble fraction after centrifugation. Therefore, the insoluble fraction was kept and resuspended in 8 M urea, 20 mM NaPi pH 7.4, 300 mM NaCl, and 10 mM imidazole. The resuspended sample was centrifuged at 125,000 x g for 40 min at 4 °C. Under denaturing conditions, FUS LC is soluble and resides in the supernatant. This supernatant was mixed with Ni-NTA resin (G Biosciences, USA) for 1 hour at 4°C. The Ni-NTA resin was then settled in a glass column and washed with the above solubilizing buffer. The bound proteins were eluted from the Ni-NTA resin with a buffer containing 8 M urea, 20 mM NaPi pH 7.4, 300 mM NaCl, and 500 mM imidazole. Purified proteins were then exchanged into a storage buffer (20 mM CAPS, pH 11) using 3K Amicon Ultra centrifugal filters (Millipore, USA). Small aliquots of the protein were flash-frozen using liquid nitrogen and stored at −80 °C.

### Protein labeling

For visualization, RGG and FUS LC were labeled with the amine-reactive Atto 488 NHS ester. The labeling reaction for RGG took place in its storage buffer (25 mM HEPES, 500 mM NaCl, pH 7.5). Dye was added to the protein in 2-fold stoichiometric excess and allowed to react for 30 min at room temperature. Labeled protein was separated from unconjugated dye using Amicon Ultra 0.5 mL centrifugal filters with MWCO of 3K. The labeling ratio was measured using UV-Vis spectroscopy. Labeled proteins were dispensed into small aliquots, flash-frozen in liquid nitrogen, and stored at −80 °C. Labeling of FUS LC followed the same process except that the labeling reaction was done in the buffer containing 50 mM HEPES at pH 7.4, and the labeled protein was then exchanged back into the storage buffer (20 mM CAPS, pH 11), divided into small aliquots, flash frozen in liquid nitrogen and stored at −80 °C.

### Suspended planar lipid membrane formation

First, lipids dissolved in chloroform were mixed in a glass vial and dried under a gentle N_2_ stream. The dried lipid film was redissolved in a mixture of hexadecane and silicone oil (1:1 v/v) to obtain a lipid/oil solution with a total lipid concentration of 3 mM. Each oil was filtered through a 0.2 μm syringe filter (Corning Inc.) before use. The lipid/oil solution was bath-sonicated for 30 min and used for experiments within several hours.

The imaging chamber was assembled by placing a PDMS gasket onto no. 1.5 glass coverslips (VWR). To make a PDMS gasket, 2.1 g of Sylgard™ 184 silicone elastomer base and 0.3 g of curing agent (Dow Corning) were thoroughly mixed and poured into a single well of a 12-well plate, where a cylindrical structure with a diameter of 8 mm was placed at the center to make a circular hole in the PDMS layer. The mixture was then incubated at 45 °C for at least 3 hours for curing. Coverslips were cleaned with 2% v/v Hellmanex III (Hellma Analytics) solution, rinsed thoroughly with deionized water, and dried under a gentle N_2_ stream before use. The coverslip was then passivated with a layer of PLL-PEG, which was synthesized as described previously.^20^ In each imaging chamber, 15 μL of PLL-PEG solution was added, followed by repetitive rinsing with the experimental buffer using a pipette after 20 min of incubation. A total of 150 μL of the aqueous buffer was then added into the PLL-PEG passivated imaging chamber. After that, 2 μL of lipid/oil solution was gently dropped and spread on the air-aqueous buffer interface. After several minutes, a hexagonal TEM grid made of gold (G150HEX Au, Gilder Grids), which was hydrophobically coated by incubating in 1-dodecanethiol solution (5 mM dissolved in ethanol) overnight before use, was gently placed on the air-oil interface using tweezers. The grid was placed there for several minutes to create a thin oil film within the grid holes. Then, the grid was submerged into the aqueous buffer using a syringe needle to place it on the PLL-PEG-coated glass surface. The thickness of the oil film decreased as the oil drained out, and spontaneous adhesion of two lipid monolayers occurred, resulting in a suspended planar lipid bilayer. Proteins were added to the aqueous buffer above the lipid bilayer.

In experiments requiring a spacer, two small strips of Scotch^®^ Magic™ Tape (3M), each a few millimeters wide, were adhered to the coverslip surface. The space between these strips was a few millimeters, slightly less than the diameter of the grid. The same procedure was followed to coat the grid with the lipid oil solution to form the membranes. When the grid was submerged in the aqueous buffer, care was taken to settle it on top of the strips of tape so that it bridged the gap between them, such that the protein solution could flow beneath the grid.

### Microscopy

A spinning disk confocal microscope (SpinSR10, Olympus) equipped with a Hamamatsu Orca Flash 4.0V3 sCMOS Digital Camera was used to visualize samples. 20X objective (1-U2M345, Olympus), 1.40 NA/40X oil immersion objective (1-UXB220, Olympus), and 1.50 NA/100X oil immersion objective (1-UXB170, Olympus) were used for visualization. Laser wavelengths of 488 and 561 nm were used for excitation.

### Image analysis

ImageJ was used for image analysis. Contrast and brightness were kept the same for time-lapse image sets for all images. For all cases, fluorescence intensity values were measured in unprocessed images.

### Mean squared displacement (MSD) analysis

Protein condensates with a defined radius were detected and tracked using the TrackMate plugin with the Laplacian of Gaussian (LoG) detector and Simple LAP tracker. MSD was calculated based on the tracked position of each condensate using a custom Matlab code.

### Coarsening dynamics analysis

The condensate area was measured using the “Analyze Particles” function in ImageJ. Growth exponent from A vs. t results was calculated using a custom Matlab code.

## Acknowledgements

This research was supported by the NIH through grant R35GM139531 (J.C.S), the NSF through grant 2218467(J.C.S), and the NSF MRSEC 1720595.

