## Supporting Information for "Transmembrane coupling accelerates the growth of liquid-like protein condensates"

**This PDF file includes:**

Figure S1
Legends for Movies S1 to S3

**Other supporting materials for this manuscript include the following:**

Movies S1 to S3

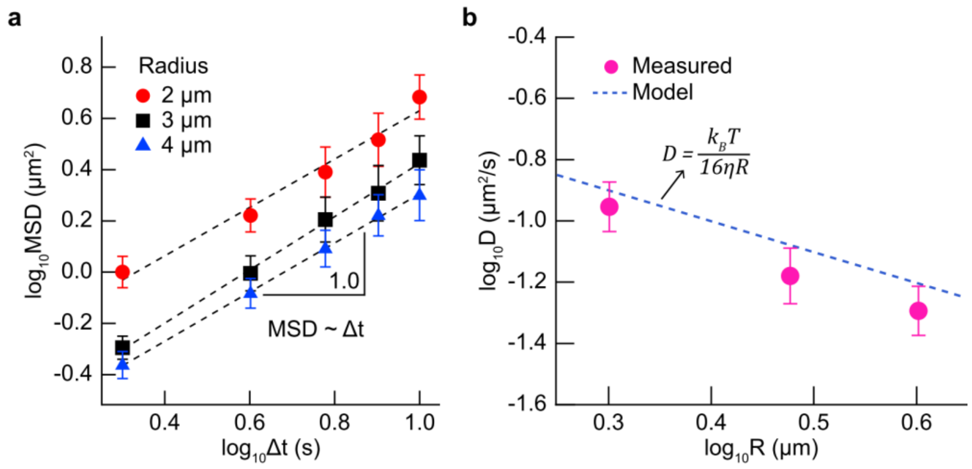


**Figure S1.** Diffusion of uncoupled FUS LC protein condensates bound on one side of the membrane exhibits Brownian motion. (a) MSDs over time lags (Δt) for FUS LC condensates with a radius of 2 µm (red circles, n = 12), 3 µm (black squares, n = 22), and 4 µm (blue triangles, n = 24). The dashed lines represent a linear fit: $\log_{10}\text{MSD} = b + \alpha \cdot \log_{10}\Delta t$. For all MSDs, the fitted slopes were 1.0 (α = 1), confirming MSD ~ Δt, a characteristic of Brownian motion. (b) Diffusivity (D) for each condensate radius (R) was obtained as $4 \cdot D = 10^b$ using the fitted results in (a): $\text{MSD} = 10^b \Delta t^\alpha$. The dashed blue line represents a model predicting D, that is, $D = k_B T / (16\eta R)$, where $k_B$ is the Boltzmann constant, T is the temperature (293 K), η is the water viscosity (1.0 cP), and R is the condensate radius. Error bars represent standard errors. Membrane composition: 80 mol% DOPC, 20 mol% DGS-Ni-NTA, and 0.5 mol% Texas Red-DHPE. Buffer: 25 mM HEPES, 300 mM NaCl, pH 7.4. 500 nM of his-FUS LC labeled with Atto 488 was used.

**Movie S1 (separate file).** A combination of representative time-lapse movies where membrane-bound RGG protein condensates are tracked according to their radii, from 1 µm to 4 µm. Tracked condensates are indicated with magenta circles. Membrane composition: 85 mol% DOPC and 15 mol% DGS-Ni-NTA. Buffer: 25 mM HEPES, 100 mM NaCl, pH 7.4. 1 µM of his-RGG labeled with Atto 488 was used. Scale bar, 10 µm.

**Movie S2 (separate file).** A combination of representative time-lapse movies highlighting a large protein composite region (left) and small coupled condensates on the bottom surface (right), described in Figure 4d. Condensates of interest are indicated with magenta circles. Scale bar, 5 µm.

**Movie S3 (separate file).** Time-lapse movie showing an increase in the extent of transmembrane condensate coupling over time, described in Figure 5a. Small uncoupled condensates on the bottom surface become continuously coupled to large condensates on the top surface, leading to an increase in the area of the brightest coupled regions. Membrane composition: 85 mol% DOPC and 15 mol% DGS-Ni-NTA. Buffer: 25 mM HEPES, 100 mM NaCl, pH 7.4. 1 µM of his-RGG labeled with Atto 488 was used. Scale bar, 5 µm.
